# Oncogenic signaling alters cell shape and mechanics to facilitate cell division under confinement

**DOI:** 10.1101/571885

**Authors:** Helen K. Matthews, Sushila Ganguli, Katarzyna Plak, Anna V. Taubenberger, Matthieu Piel, Jochen Guck, Buzz Baum

**Affiliations:** MRC Laboratory for Molecular Cell Biology, University College London, Gower Street, WC1E 6BT, UK; The Francis Crick Institute, 1 Midland Road, London NW1 1AT, UK; Biotechnology Center, Technische Universität Dresden, Tatzberg 47/49, 01307 Dresden, Germany; Institut Curie, PSL Research University, CNRS, UMR 144, Paris 75005, France; Institute for the Physics of Living Systems, University College London, London, WC1E 6BT, UK

## Abstract

When cells enter mitosis, they become spherical and mechanically stiffen. We used MCF10A cell lines as a model system in which to investigate the effect of induced oncogene expression on mitotic entry. We find that activation of oncogenic Ras^V12^, for as little as five hours, changes the way cells divide. Ras^V12^-dependent activation of the MEK-ERK signalling cascade alters acto-myosin contractility to enhance mitotic rounding. Ras^V12^ also affects cell mechanics, so that Ras^V12^ expressing cells are softer in interphase but stiffen more upon entry into mitosis. As a consequence, Ras^V12^ expression augments the ability of cells to round up and divide faithfully when confined underneath a stiff hydrogel. Conversely, inhibition of the Ras-ERK pathway reduces mitotic rounding under confinement, resulting in chromosome segregation defects. These data suggest a novel mechanism by which oncogenic Ras-ERK signalling can aid division in stiff environments like those found in tumours.

## Introduction

Animal cells undergo profound changes in both cell shape and mechanics at the start of mitosis. In tissue culture, adherent spread cells retract their margins in early mitosis as they round up to become spherical^1^ - a process driven by a combination of substrate de-attachment^2^, acto-myosin re-modelling^3–5^ and osmotic swelling^6–8^. At the same time, cells become stiffer^3,4,9^. This change in cell mechanics requires the re-modelling of actin filaments into a thin network at the cell cortex^10^ and is essential for cells to divide in a stiff gel that mimics a tissue environment^11^. Because flattened cells lack the 3-dimensional space required to assemble a bipolar spindle and capture chromosomes^12^, physical confinement results in multiple defects in spindle formation and chromosome segregation^13^.

While almost all proliferating animal cells undergo a degree of mitotic rounding, different cell types exhibit striking differences in the extent to which they round^1,12^. In this context, we previously noted that cancer cell lines tend to round up more than many non-transformed cells^2^. There are two likely explanations for this. First, the ability of a cell to successfully build a spindle in a flattened state depends on centrosome number and DNA content^12,13^. This is important since cancer cells tend to have more chromosomes and centrosomes than non-transformed cells. HeLa cells for example have close to three times the normal number of chromosomes^14^. In line with this, cancer cells suffer greater mitotic defects than non-transformed cells when rounding is limited by mechanical constraints^12,13^. Second, while normal cells divide in a defined tissue niche where the mechanical and physical environment is tightly regulated, to survive cancer cells must be able to divide in a wide range of environments including a crowded primary tumour, in the circulatory system^15^ and at metastatic sites; all of which have biochemical and mechanical properties that are very different to those in the original tissue. While the nature of the genetic changes that enable cancer cells to divide in different environments is not known, we have previously shown that the acto-myosin cytoskeleton controls mitotic rounding^3,4,13,16^. This led us to put forward the hypothesis that regulators of the acto-myosin cortex may be co-opted by cancer cells to enable them to successfully divide in different environments^17^. Indeed, many of the proteins required for mitotic rounding, such as Ect2 and Ezrin are up-regulated in cancer^18,19^. However, it is difficult to directly compare mitotic rounding and cell division in normal and cancer cells, not least because of the large number of changes that cells accumulate during transformation. Therefore, as an experimental system in which to study how transformation influences mitotic rounding, we chose to induce the expression of oncogenes in a non-transformed diploid epithelial cell line: MCF10A cells. Remarkably, in this model system, five hours of expression of a single oncogene, Ras^V12^, was sufficient to profoundly alter mitotic cell shape dynamics and mechanics - mirroring the division changes seen in cells with sustained, long-term Ras^V12^ overexpression. At the same time, Ras^V12^ activation was able to induce mechanical changes that improved the ability of cells to round up and faithfully divide underneath gels designed to mimic a stiff extracellular environment. These data show how oncogenes can aid the high-fidelity division of cancer cells to extend their ability to proliferate across a range of environments - a likely pre-requisite for cancer progression.

## Results

### Activation of oncogenic Ras alters mitotic cell geometry

To investigate the impact of oncogenic signaling on mitotic cell shape and mechanics we used the non-transformed human epithelial cell line (MCF10A) as a model system ^20^. Oncogenic h-Ras^G12V^ (hereafter called Ras^V12^) was activated in these cells by constitutive over-expression or using an inducible system in which Ras^V12^ can be rapidly activated and stabilized following addition of 4-OH-tamoxifen^21^. Over-expression of full length or ER-fused Ras and phosphorylation of its downstream target, ERK, were confirmed for both lines using Western blotting (Figure 1A). To determine how cells change shape upon entry into mitosis, brightfield time-lapse microscopy was used in each case to follow unlabeled asynchronous populations of cells as they divided (Figure 1B). Cell spread area and aspect ratio were measured as cells underwent rounding. When we quantified cell length (Feret diameter) and aspect ratio 15 minutes before mitotic entry and in metaphase (5 minutes before anaphase elongation) for each population (Figure 1C), it became clear that a brief induction of Ras^V12^ expression was sufficient to enhance mitotic rounding (Figure 1C, S1A). Strikingly, this effect was similar to that seen in cell lines constitutively expressing Ras^V12^, as a model of long-term Ras activation (Figure 1C). A similar effect was observed following inducible activation of k-Ras^G12V^ (Figure S1B). Since activation of Ras^V12^ likely also affects the stability of cell/cell junctions, we asked how this effect is influenced by the presence of neighbouring cells. We observed that MCF10A cells rounded more upon entry into mitosis if cultured at low density, compared to cells surrounded by neighbours in the middle of a monolayer. However, shortterm Ras^V12^ induction enhanced mitotic rounding in single cells and cells within a monolayer (Figure 1D), which then rounded to a similar extent. This suggests that Ras activation, for as little as 5 hours, alters cell/cell junctions and monolayer integrity, even though these cells continue to express E-cadherin (Figure 1A). Importantly, the difference in the rate of rounding observed in single cells suggests that Ras^V12^ also induces intrinsic changes to the way that cells change shape in mitosis. Since the aim of this work was to investigate these intrinsic differences, all further experiments were carried out with cells plated at low density to exclude effects of cell/cell interactions on cell shape.

**Figure 1.**
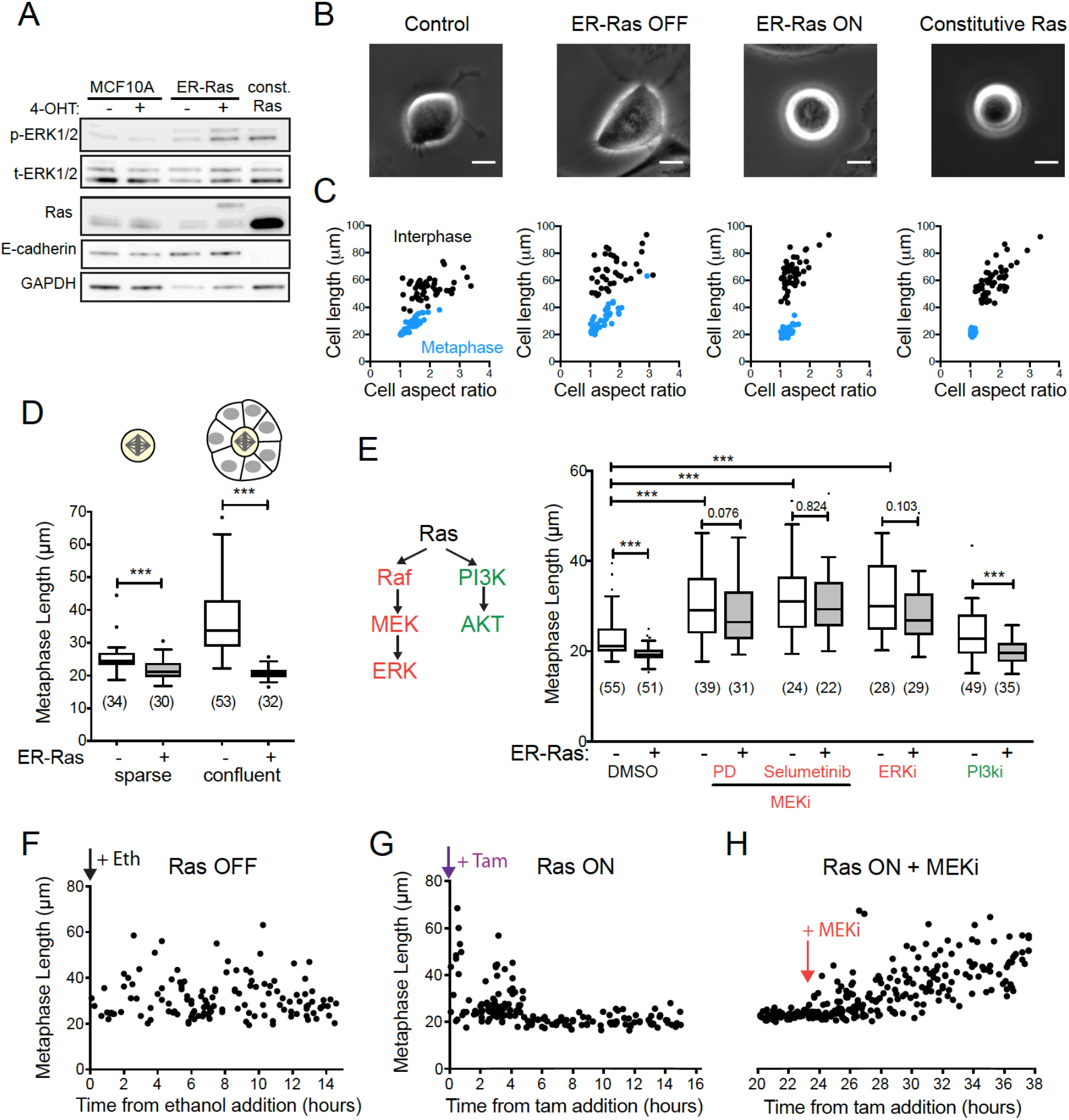
Ras/MEK/ERK signaling controls cell shape in mitosis. (A) Western blot showing levels of ERK1/2 phosphorylation and total ERK1/2, Ras and E-cadherin expression from MCF10A, MCF10A ER-hRas^V12^ and MCF10A+hRas^v12^ cells. MCF10A and ER-hRas^V12^ cells were treated with ethanol (-) or 4-OH-tamoxifen (+) for 7 hours before lysis. (B) Representative phase contrast images of the different cell types in metaphase. Metaphase is taken as 5 minutes before anaphase elongation or furrowing is first observed. Scale bars are 10 μm. (C) Scatter plots of cell length (Feret’s diameter) and aspect ratio for the different cell types in interphase (black, taken as 15 minutes before nuclear envelope breakdown) and metaphase (blue, 5 minutes before anaphase). Cells were imaged every 5 minutes for 15 hours using phase contrast microscopy and the shapes recorded for every cell division. For ER-hRas^V12^ cells, cells were analysed during the period of 5 – 15 hours post ethanol or 4-OH-tamoxifen addition. Short and long term Ras activation had a significant effect on mitotic cell length (control vs constitutive Ras***, ER-Ras OFF versus ER-Ras ON***) and aspect ratio (control vs constitutive Ras***, ER-Ras OFF versus ER-Ras ON p = 0.0046**) n = 30 cells per condition. (D) Box plot showing metaphase length for MCF10A ER-hRas^V12^ following 5 - 15 hours ethanol or 4-OH-tamoxifen treatment comparing cells plated in sparse (defined as single cells or cells with <2 neighbours) or confluent (cells surrounded by other cells on all sides) conditions. (E) Box plot showing metaphase length for ER-hRas^V12^ following 5 - 15 hours ethanol or 4-OH-tamoxifen treatment alongside addition of DMSO or the following small molecule inhibitors: 2 μM PD 184352, 10 μM Selumetinib (MEK inhibitors), 10 μM GDC-0994 (ERKi) and 2 μM ZSTK474 (Pi3Ki). (F-H) Plot of metaphase length of individual ER-hRas^V12^ cells dividing against time after ethanol (F) or 4-OH-tamoxifen (G) addition and following addition of 10 μM Selumetinib (H).

### Ras signals through the MEK/ERK pathway to control mitotic cell shape

One of the main effectors of Ras activation is the MEK-ERK signaling cascade and we observed increased ERK phosphorylation following both inducible and long-term Ras activation (Figure 1A). To test whether this pathway controls mitotic shape, we inhibited different elements of the pathway in cells with and without Ras^V12^ induction. Inhibition of MEK or ERK (but not PI3K) abolished the difference in mitotic cell shape seen following Ras activation (Figure 1E). Furthermore, MEK or ERK inhibition significantly increased metaphase cell length in both control MCF10A cells (Figure 1E, Figure S1D) and in constitutive Ras^V12^-expressing cells (Figure S1E). This reveals a previously unidentified role for the Ras-ERK signalling cascade in the control of mitotic rounding. We examined the timing of these Ras-ERK dependent changes in mitotic cell shape and found that the change in metaphase length began around 5 hours after Ras^V12^ activation (Figure 1F, G) and was reversed within 30 minutes of treatment with a MEK inhibitor (Figure 1H). These data demonstrate that the Ras-ERK signaling pathway plays a role in controlling the shape of cells in mitosis. More surprisingly, these changes emerge over a timescale of minutes/hours and are not simply a long-term consequence of oncogenic transformation.

### Ras-ERK signaling alters cell contractility in mitosis

To determine how the dynamics of mitotic rounding was altered following Ras activation, we constructed MCF10A lines that stably express LifeAct-GFP to visualise actomyosin dynamics through mitosis (Figure 2). By following the shape changes of individual cells, we observed that mitotic rounding begins 5-10 minutes before nuclear envelope breakdown (NEB) (Figure 2A, B) as previously observed in HeLa cells ^3^. Rounding was initiated at a similar time in cells with or without Ras^V12^ induction, but Ras^V12^ expressing cells rounded at a faster rate, especially before NEB (Figure 2B). This was the case for cells induced to express Ras^V12^ and for cells overexpressing constitutive Ras^V12^ (Figure 2C). Thus, cells expressing oncogenic Ras^V12^ were almost completely round by pro-metaphase while control cells or cells treated with a MEK inhibitor continued to round up throughout mitosis until they entered anaphase and divided. This was reflected in a significant difference in rounding rates, with Ras cells rounding faster than controls; a difference that was abolished by treatment with a MEK inhibitor (Figure 2D). These differences resulted in metaphase cells that were narrower and taller following Ras activation and conversely, wider and flatter following pathway inhibition using a MEKi (Figure 2E, Figure S2A).

**Figure 2.**
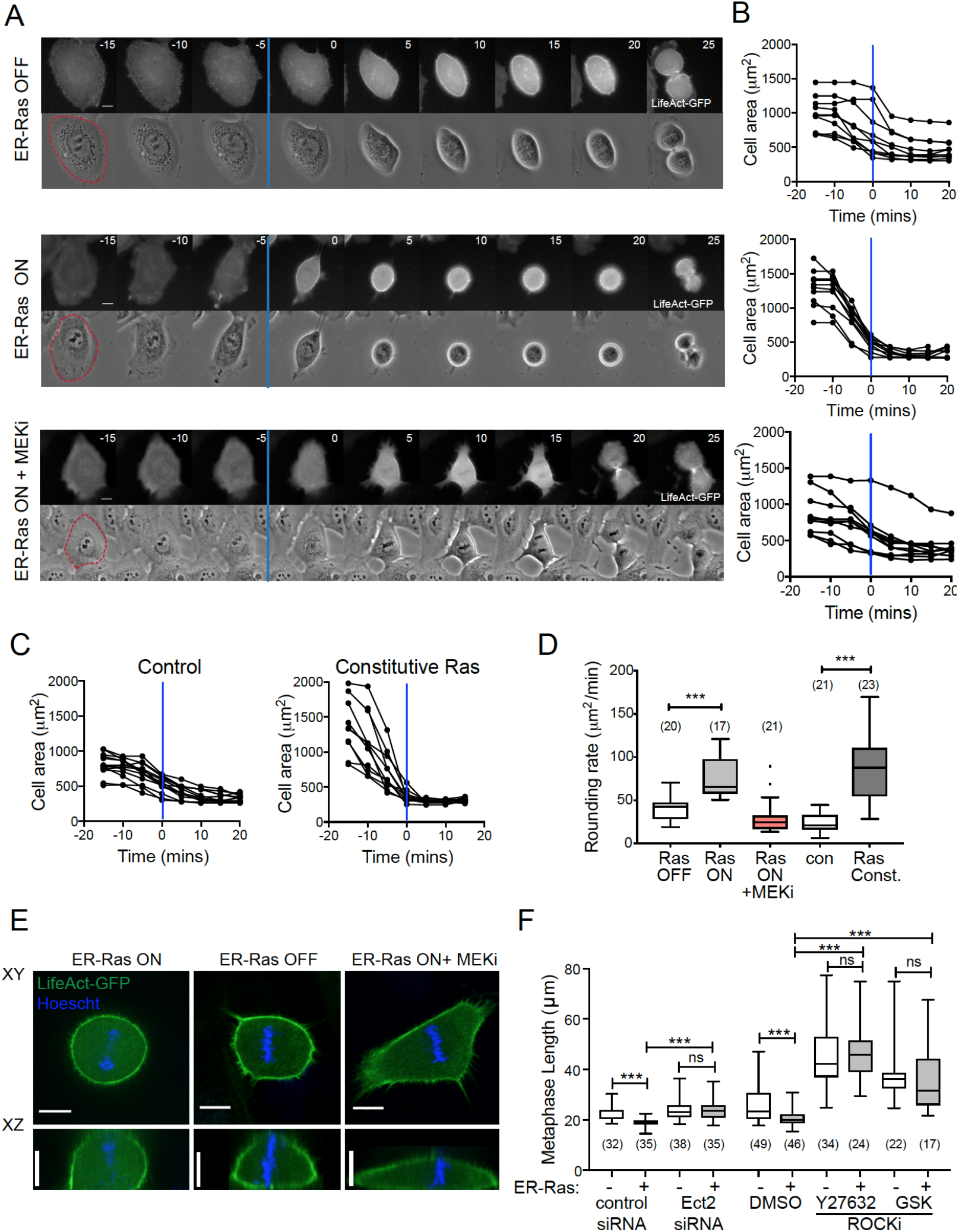
Ras/MEK/ERK signaling alters contractility in early mitosis. (A) Representative montage time-lapse images of MCF10A ER-hRas^V12^ cells labeled with LifeAct-GFP following 8 hour treatment with ethanol, 4-OH-tamoxifen or 4-OH-tamoxifen + 2 μM PD 184352 (MEKi). Time in minutes is aligned so that t = 0 is frame after NEB (blue line). Scale bars are 10 μm (B) Quantification of cell area for 8 cells from conditions in A entering mitosis, aligned so that t = 0 is NEB (blue lines). Measurements were taken from time-lapse microscopy of cells 5 -15 hours after treatment. (C) Quantification of cell area for control MCF10A cells and MCF10A+hRas^v12^ entering mitosis, aligned so that t = 0 is NEB (blue lines). (D) Box plot showing mean rate of area decrease during mitotic rounding for conditions shown in A-C. (E) XY and XZ confocal slices of metaphase MCF10A ER-hRas^V12^ - LifeAct-GFP cells following 7 hours treatment with ethanol, 4-OH-tamoxifen or 4-OH-tamoxifen + 2 μM PD 184352 (MEKi). DNA was visualized by treatment with Hoechst for 10 minutes prior to imaging. Scale bars are 10 μm. (F) Box plot showing metaphase cell length for ER-hRas^V12^ following 5-15 hours ethanol or 4-OH-tamoxifen treatment and treated with control or Ect2 siRNA, or DMSO, 25 μM Y-27632 or 2 μM GSK 269962 (ROCK inhibitors).

Since Ras activation did not affect cell volume at mitosis (Figure S2B), mitotic swelling ^6–8^ appeared unaltered. Instead, it is likely that the Ras-ERK pathway functions to alter acto-myosin contractility. Actin rearrangements in early mitosis are driven by the activation and re-localisation of the RhoGEF, Ect2 ^3^, which activates RhoA ^5^ and its downstream effector Rho kinase (ROCK) to contract the cell edge. We found that inhibition of this pathway though siRNA knockdown of Ect2 or small molecular inhibitors of ROCK abolished the effect of Ras activation on cell shape (Figure 2F). Taken together, these data show that activation of the oncogenic Ras/MEK/ERK signalling pathway is sufficient to induce changes in mitotic cell shape in epithelial cells over both short and long timescales and that these shape changes require Ect2 - mediated changes in acto-myosin contractility.

### Ras activation alters cell mechanics

Acto-myosin rearrangements at mitotic entry not only alter cell shape but also their mechanical properties. Thus, mitotic cells are significantly stiffer than interphase cells ^3,4,9^. This led us to compare the mechanical properties of normal MCF10A cells and Ras^V12^ over-expressing cells using two complementary techniques. First, we measured the apparent elastic modulus of individual cells using atomic force microscopy (AFM). To control for the effect of cell shape on mechanics, cells were first removed from the substrate using a brief treatment with Trypsin-EDTA and then allowed to re-attach but not spread on the substrate. In this way, we could directly compare the stiffness of interphase and mitotic cells with the same shape (Figure 3A). Under these conditions, Ras^V12^ expressing cells were significantly softer in interphase than controls, as previously reported for rounded cells ^22^. At the same time, Ras^V12^ expressing cells were marginally but significantly stiffer than control cells in mitosis (Figure 3A). As a consequence, Ras^V12^ cells underwent a much greater fold change increase in their elastic modulus upon mitotic entry than controls (4.21x mitotic increase for Ras^V12^ cells, 2.23x mitotic increase for controls). Similar results were obtained using real-time deformability cytometry (RT-DC)^23^ as a complementary approach to measure cell stiffness (Figure 3B). Whereas AFM probes the cortical mechanics of cells sitting on a surface, RT-DC is a high throughput method (>2000 cells per experiment) to quantify the global mechanical properties of cells in suspension. Using RT-DC, we confirmed that Ras^V12^-expressing cells were significantly softer than their non-transformed counterparts in interphase but were similar in stiffness in mitosis (Figure 3B). These data show that long-term Ras activation causes cells to stiffen more at mitosis in a manner that is independent of their shape.

**Figure 3.**
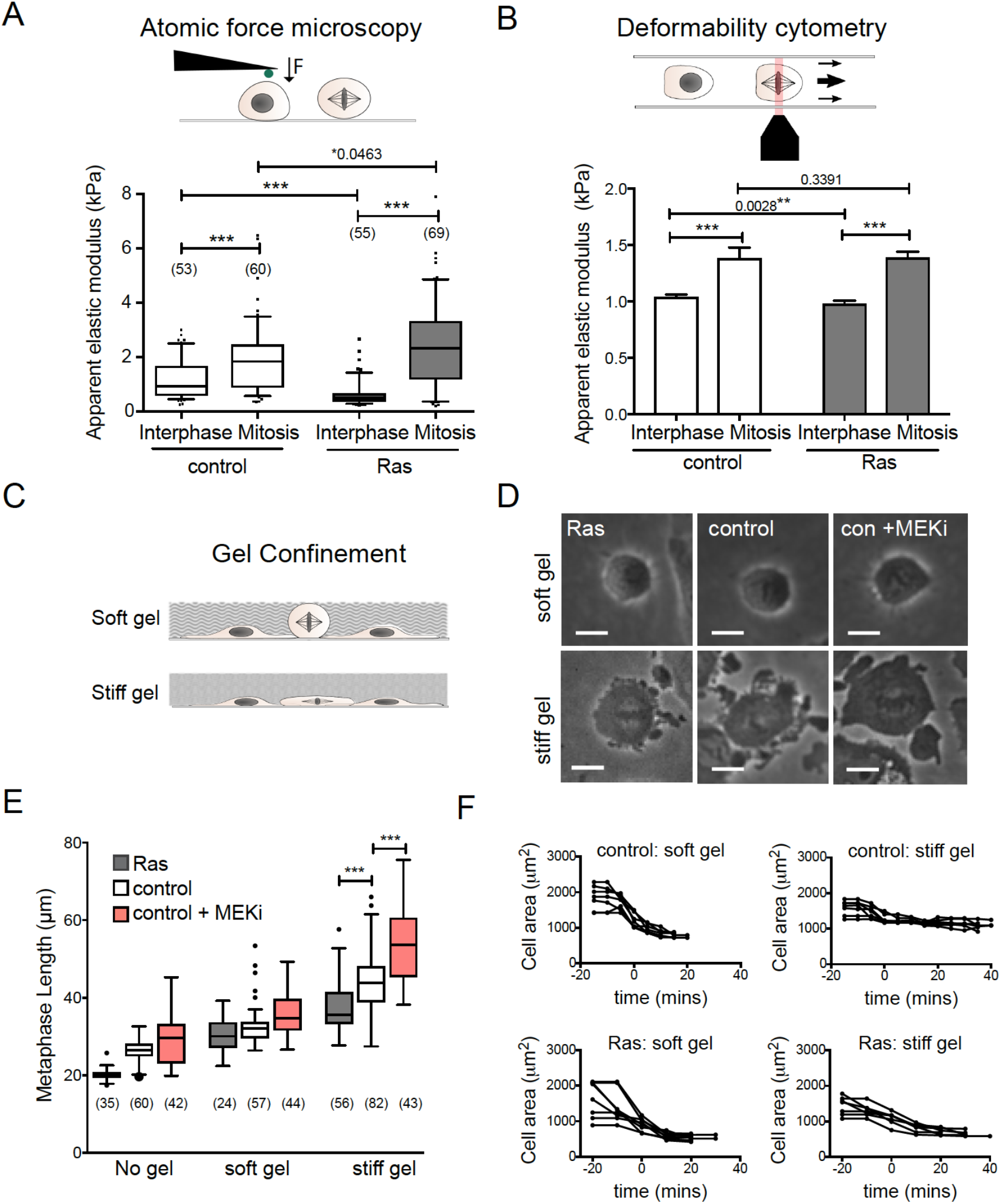
Ras activation alters cell mechanical properties. (A) Schematic of atomic force microscopy (AFM) set-up. Box plot showing the apparent elastic modulus for rounded control MCF10A cells and MCF10A+hRas^v12^ in interphase and mitosis (arrested in pro-metaphase by STLC treatment) as determined by AFM. Graph shows 1 representative example from 2 independent experiments. (B) Schematic of RT-DC set-up. Bar chart shows average apparent elastic modulus values calculated for interphase and mitotic cells from 10 independent experiments (n > 15000 cells). Due to large sample size a linear mixed model was used to calculate the statistical significance of the results. (C) Schematic of cells dividing in confinement under compliant (~5 kPa) and stiff (~ 30 kPa) polyacrylamide gels. (D) Phase contrast images of MCF10A+hRas^v12^, control MCF10A and MCF10A + 2 μM PD 184352 cells in metaphase under soft and stiff gels. Scale bars are 10 μm. (E) Box plot showing metaphase cell length for MCF10A+hRas^v12^, control MCF10A and MCF10A + 2 μM PD 184352 without confinement and under soft and stiff gels. Data comes from timelapse movies for 15 hours after confinement onset. The MEKi was added immediately before confinement and mitoses were analysed 5 - 15 hours post addition/confinement. (F) Graphs showing cell area for control MCF10A cells and MCF10A+hRas^v12^ entering mitosis under soft and stiff gels, aligned so that t = 0 is NEB.

### Ras activation promotes mitotic rounding under confinement

Next, we sought to investigate how these changes in the rate and extent of rounding and the changes in cell mechanics induced by Ras^V12^ influence cell division under confinement. To do this, we confined cells under polyacrylamide hydrogels of different stiffness^24^; a compliant gel of around ~5 kPa and a stiff gel of around ~30 kPa (Figure 3C) and filmed cells as they proceeded through mitosis in these confined conditions. Under the soft gel, cells rounded up normally upon entry into mitosis. By contrast, under the stiffer gel, the ability of cells to round was severely limited (Figure 3D, E). Strikingly, however, the expression of Ras^V12^ improved the ability of cells to round up under the stiff gel – leading to a significant reduction in metaphase length compared to controls cells (Figure 3E, mean length for controls was 44.2 ± 7.6 μm, for Ras^V12^ cells 37.5 ± 6.2 μm). Conversely, cells treated with a MEK inhibitor remained even flatter when confined under the stiff gel (Figure 3E, mean length 55.1 ± 12.5). This was reflected in a change in the dynamics of rounding. More strikingly still, while control cells grown under a stiff gel were unable to retract their margins, this inhibition could be relieved by the expression of oncogenic Ras^V12^ (Figure 3F). Thus, the alterations in cell shape and contractility induced by Ras^V12^ over-expression allow cells to round up better in confined conditions, suggesting that they are able to exert more force to deform the overlying gel as they enter mitosis.

### Ras-induced enhanced mitotic rounding reduces mitotic defects under confinement

Limiting mitotic rounding by confinement results in multiple mitotic defects ^13^. This led us to test whether the enhanced ability of Ras^V12^ cells to round up under a stiff gel provided any advantage for cell division in these conditions. Firstly we measured the time taken by cells to pass through mitosis (nuclear envelope breakdown to anaphase) (Figure 4A). This serves as a good proxy for spindle defects, as anaphase is only able to proceed once all chromosomes are attached to the spindle to satisfy the spindle assembly checkpoint^25^. When cells were confined under a soft gel, we did not observe any change in the timing of mitosis. However, increasing gel stiffness resulted in a prolonged mitosis in all conditions (Figure 4A). Ras^V12^ cells, which were better able to round up under the gel (Figure 3E, F), were significantly faster at progressing through mitosis compared to normal MCF10A cells (Figure 4A). The delays in mitotic progression were associated with profound differences in spindle integrity under stiff gels. Under the stiff gel, spindles often fractured to produce a tri-polar spindle^13^, resulting in a cell dividing into three rather than two daughter cells. However the incidence of spindle fracturing was decreased in Ras^V12^ expressing cells and increased following treatment with a MEK inhibitor (Figure 4B). We also observed signs of cortex instability such as extreme blebs in mitotic cells under stiff gels. These blebs were often longer than the cell itself, and were sometimes seen breaking off from the cell body. Again, this type of extreme blebbing was reduced in cells expressing Ras^V12^ compared to normal MCF10A and cells in which MEK had been inhibited (Figure 4C). This suggests that the ability of Ras^V12^ cells to round up under the stiff gel is able to limit some of the mitotic defects induced by confinement. To test this, we blocked acto-myosin contractility a using a ROCK inhibitor. Under these conditions, mitotic rounding was limited under the soft gel as well as the stiff (Figure 4D) leading to a prolonged mitosis under both gel types (Figure 4E) but not in unconfined cells (Figure S2C, D). Crucially, this also abolished any difference between Ras^V12^ expressing and control cells (Figure 4D, E). This demonstrates that the acto-myosin cytoskeleton is essential to generate the pushing force that enables Ras^V12^ cells to round up and successfully divide in confined conditions.

**Figure 4.**
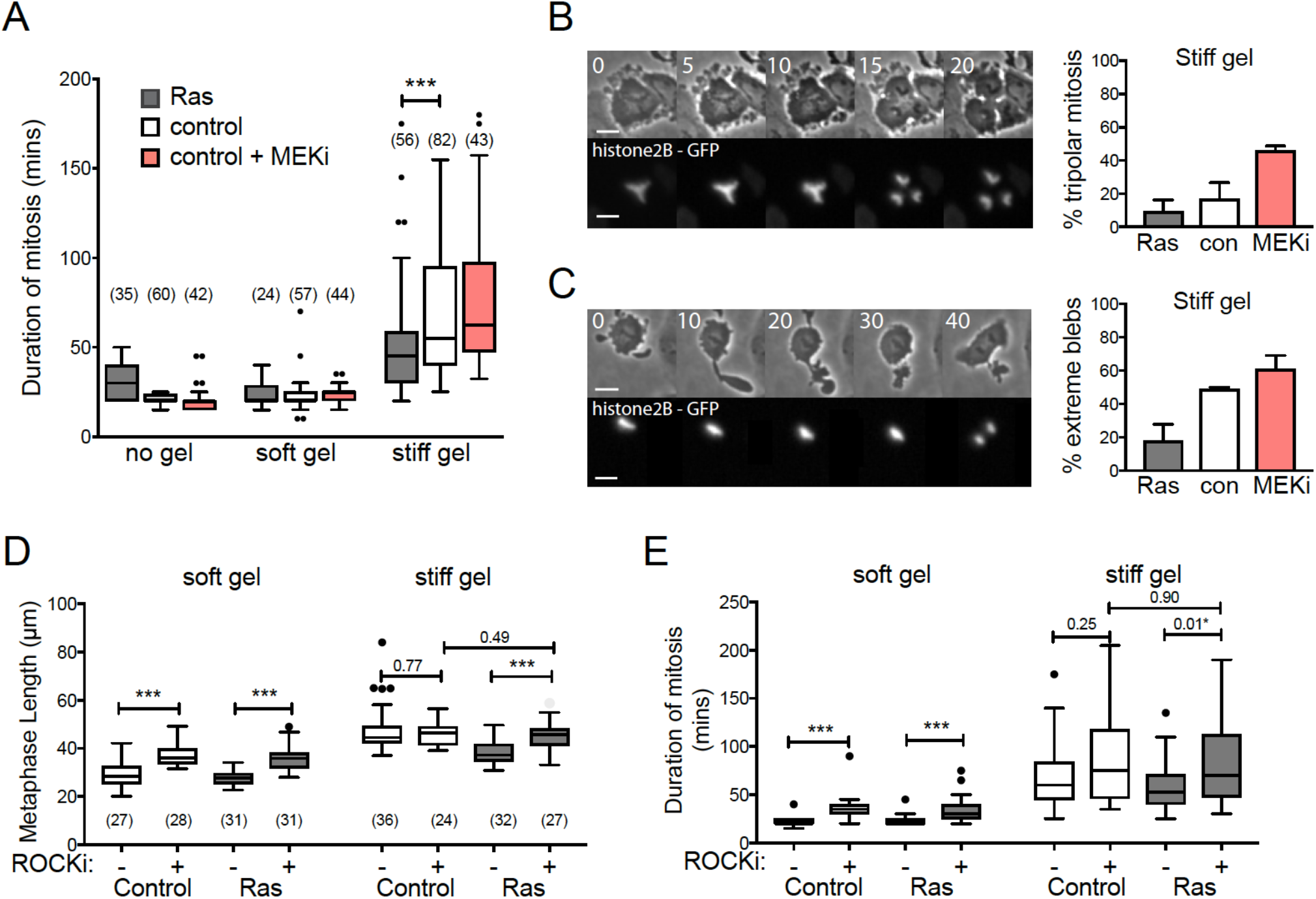
Ras activation facilitates accurate and timing cell division in confinement. (A) Box plot showing the time taken for cells to progress through mitosis (from NEB until anaphase) for MCF10A+hRas^v12^, control MCF10A and MCF10A + 2 μM PD 184352 without confinement and under soft and stiff gels. Data comes from timelapse movies for 15 hours after confinement onset. MEKi was added immediately before confinement and mitoses were analysed 5 - 15 hours post addition/confinement. (B) Time-lapse montage showing example of tri-polar mitosis and graph showing % of cells that undergo tri-polar mitosis underneath a stiff gel. N = 3 independent experiments. (C) Time-lapse montage showing an example of extreme blebbing and graph showing % of cells that undergo extreme blebbing underneath a stiff gel. Extreme blebbing was defined as any mitotic cells having a bleb equal to or longer than cell diameter. N = 3 independent experiments. Note that no tri-polar mitosis or extreme blebbing was observed in any condition under soft gels or without gels. Times are in minutes, scale bars are 20 μm. (D, E) Box plot of metaphase length (D) and mitotic duration (time from NEB to anaphase) (E) for control MCF10A and MCF10A+hRas^v12^ cells under soft and stiff gels, treated with DMSO or 25 μM Y-27632 (ROCKi).

## Discussion

Activation of oncogenic Ras is a driver mutation in many cancer types and leads to multiple downstream events that promote transformation^26^. Here we have discovered a novel role for Ras and the downstream MEK-ERK pathway in directly controlling the geometry and mechanics and rounding force generation of cells during mitosis. Using an inducible cell line, we have shown that oncogenic Ras^V12^ activation, for as little as five hours, makes cells more spherical at metaphase. These geometric changes are also observed following long-term Ras^V12^ overexpression, along with corresponding changes in cell mechanics and the ability to apply force under a gel. This enables Ras^V12^ cells to round up even when confined underneath a stiff hydrogel that limits the rounding of control cells and to more effectively build a mitotic spindle and divide in confined conditions. This hints at a wider role for oncogenic signaling in directly controlling the mechanics of cell division to facilitate division in stiff or confined environments.

Activation of mutated Ras leads to a cascade of events over different timescales, from the phosphorylation of signaling proteins such as ERK (minutes)^21^ to changes to cytoskeletal architecture (hours) to cell state changes such as epithelial-to-mesenchymal transition (days). Our observation that mitotic shape changes begin five hours after Ras activation suggests that this is not due to longterm mutations accumulating over many days. Given the timeframe (5 hours for induction and 30 minutes following ERK inhibition), this may require alterations in gene expression, likely for genes that regulate the actin cytoskeleton, whose function our data show is required for the effect. This is in line with the well-documented role of the Ras-ERK pathway in controlling actin filament organization in interphase to promote cell motility and invasion^27,28^. Ras also affects interphase cell mechanics; inducible activation softens or stiffens cells in a substrate-dependent manner^22,29^ while long-term overexpression softens interphase cells^30^, as we have observed here. Ras also regulates acto-myosin rearrangements at mitotic exit to determine how daughter cells re-spread after division ^31^. In a similar manner, inducible activation of oncogenic Src, which also stimulates ERK signaling, has been shown to change the expression levels of multiple actin binding proteins after several hours^32^. While the targets of Ras-ERK signalling in controlling mitotic rounding remain to be elucidated, mitotic rounding is triggered by RhoA ^5^, following activation by its GEF, Ect2^3^. Ras has been shown to modulate the activity of RhoA^33–36^, and its function in controlling mitotic shape likely also requires RhoA, since inhibition of Ect2 or ROCK block this effect (Figure 2).

The function of these oncogene-induced changes to mitotic geometry and mechanics become most apparent when cells are challenged to divide underneath a stiff gel. This illustrates the importance of environmental context on cell division; in 2D tissue culture, mitotic rounding primarily requires loss of substrate adhesion. However, under confinement, the cell’s ability to generate force to push against its environment is crucial^13^. This is also the case in 3D culture, where mitotic rounding generates force in all directions^11^, similar to cells in the tissue context. Environment context is also important for the fidelity of cell division: recent work has shown that primary epithelial cells cultured in 3D suffer fewer chromosome segregation defects than in 2D^37^. Conversely, cell culture in a stiff 3D gel that limits anaphase elongation also induces chromosomal segregation defects ^11^. This is especially relevant to tumours where profound changes to tissue integrity and mechanical properties^38,39^ likely impact the cell division process and thus drive chromosomal instability. Here we have shown that oncogenic Ras-ERK signaling enables cells to overcome mechanical constraints in their local environment that would otherwise impair their ability to divide. It is perhaps surprising then, that in this context, Ras activation is protective against chromosomal instability rather than promoting it. This may be because confinement underneath a very stiff gel (~30 kPa) is an extreme situation in which normal cells would never normally need to divide. To survive and proliferate, cancer cells, on the other hand, must be both robust to environmental challenges and insensitive to any defects in chromosomal segregation that these may cause. Thus, Ras^V12^ cells are better able to faithfully divide under stiff gels, but still suffer minor defects in chromosomal segregation (Figure 4), which may be better tolerated in the long term. Thus, the acquisition of oncogenic Ras ^V12^ is likely to provide cells with the ability to divide under a wider range of conditions and to accommodate changes in ploidy that require a larger spindle. In this way Ras^V12^ may set the stage for future evolution of the tumour.

In summary, oncogenic Ras-ERK signaling has been shown to induce multiple changes in cell biology that enable cancer cells to grow, divide and evolve in different environments and that enable them to tolerate large-scale changes in cancer genome. Here we show that this includes changes to the mechanical process of cell division itself.

## Materials & Methods

### Cell lines & culture

Cells were cultured in DMEM F-12 Glutamax, with 5% Horse serum (Invitrogen), 20ng/ml EGF (Peprotech), 0.5mg/ml Hydrocortisone (Sigma), 100ng/ml Cholera toxin (Sigma), 10μg/ml Insulin (Sigma), 1% Penstrep (Gibco). Cell lines used were MCF10A (Horizon), MCF10A+ constitutive hRas^G12V^ (gift from Susana Godinho) and the inducible lines MCF10A-ER:hRas^G12V^ and MCF10A-ER:kRas^G12V^(gifts from Julian Downward)^21^. Ras was activated in inducible lines by addition of 100nM 4-OH-tamoxifen (Sigma). LifeAct-GFP labeled lines were produced by infection with puromycin resistant lentivirus (rLV-Ubi-LifeAct-GFP2, Ibidi 60141). GFP positive cells were sorted using flow cytometry to produce a polyclonal stable pool.

### Small molecule inhibitors & siRNA

The following small molecule inhibitors were used in this study: MEK inhibitors: 2 μM PD 184352 (Cell Signaling 12147) and 10 μM Selumetinib (Selleckchem S1008), ERK inhibitor: 10 μM GDC-0994 (Selleckchem S755403), PI3k inhibitor: 2 μM ZSTK474 (Selleckchem S1072), ROCK inhibitors: 25 μM Y-27632 (Sigma Y0503) and 2 μM GSK 269962 (Tocris 4009). Details of treatment times are described in figure legends. For Ect2 knockdown, Hs_ECT2_6 (ATGACGCATATTAATGAGGAT-Qiagen SI03049249) was used as previously described^2,3^ and compared to the AllStars negative control siRNA (Qiagen 1027280). Cells were transfected using Lipfectamine RNAimax (Invitrogen 13778-075) and imaged from 24 hours post transfection.

### Live cell imaging

Cells were plated in fibronectin-coated, glass-bottomed plates (Mattek) 24 hours before imaging. Widefield timelapse imaging was carried out using a Nikon Ti inverted microscope or a Zeiss Axiovert 200M microscope at 5 minute timepoints using a 20x or 40x objective. Live confocal imaging was carried out on a Nikon TiE inverted stand attached to a Yokogawa CSU-X1 spinning disc scan head using a 100x objective. Cell shape descriptors were measured using Fiji following manual segmentation of cell body.

### Atomic force microscopy

To enrich for mitotic cells, cells were treated for 12-15 hours with 5 μM STLC (Sigma 164739). Then, the fraction containing loose mitotic cells was washed off and recombined with interphase cells that had been detached using Accutase (PAA laboratories). Cells were stained with Hoechst for 10 minutes at 37 °C, washed twice and resuspended in CO_2_-independent medium (LifeTechnologies). Cells were seeded into glass bottom dishes (FluoroDish−, WPI) and probed once they were adhered but not spread. During the AFM indentation experiment, interphase and mitotic cells were distinguished using epifluorescence images of the Hoechst stained cells. For AFM indentation measurements, a Nanowizard I equipped with a CellHesion module (JPK Instruments) was used. Arrow-T1 cantilevers (Nanoworld) were modified with a polystyrene beads (radius 2.5μm, microparticles GmbH) with the aid of epoxy glue to obtain a well-defined indenter geometry and decrease local strain during indentation. Cantilevers were calibrated prior to experiments using built-in procedures of the SPM software (JPK Instruments). The bead was lowered at a defined speed (10μm/sec) onto the cell surface. After reaching the setpoint of 2nN, the cantilever was retracted. During the force-distance cycle, the force was recorded for each piezo position. The resulting force-distance curves were transformed into force-versus-tip sample separation curves ^40^ and fitted with the Hertz/Sneddon model for a spherical indenter ^41^ using the JPK data processing software (JPK DP, JPK Instruments). A Poisson ratio of 0.5 was used for the calculation of the apparent Young’s modulus. Since rounded cells were probed, apparent Young’s moduli were corrected according to Glaubitz et al ^42^ using the average cell diameters of each cell population that had been determined in phase contrast images using FIJI. Each cell was probed three times, and an average apparent Young’s modulus calculated. As a control, cells that had not been treated with STLC nor Hoechst were probed and phase contrast was used to identify mitotic cells.

### Real-time Deformability Cytometry

Mitotic cells were collected after 5 hours of STLC treatment (5μM) using the mitotic shake-off method. Interphase cells were detached by treatment with Accutase (PAA laboratories) for 15 minutes. The cells were washed with PBS and resuspended in 0.5% methylcellulose buffer before loading in the RT-DC setup as described previously ^43^. Cells were deformed by flowing them through 30μm x 30μm channels with a speed of 0.16μl/sec or 0.32μl/sec and the deformation was recorded using ShapeIn software (Zell Mechanik Dresden). Data for over 2000 cells was recorded for each of 10 biological replicates of the experiment. Apparent elastic modulus was calculated using ShapeOut Software (Zell Mechanik Dresden) based on a numerical simulation model described previously ^44^. The results from 7 independent experiments were analysed using a Linear Mixed Model method integrated in ShapeOut Software (Zell Mechanik Dresden)^45^.

### Polyacrylamide gel confinement

Polyacrylamide gels were polymerized on 18mm glass coverslips. Coverslips were first functionalized by plasma cleaning for 30s (Diener Femto), followed by incubation with a solution of 0.3 % Bind-Silane (Sigma M6514)/5% acetic acid in ethanol. Coverslips were then rinsed with ethanol and dried with compressed air. Polyacrylamide gel solutions were made up as 1 mL solutions in PBS as follows: Stiff gels (~30 kPa): 250 μL acrylamide 40% (Sigma), 100 μL bisacrylamide 2% (Fisher Scientific), 10 μL APS (10% in water, Sigma), 1 μL TEMED (Sigma). Soft gels (~5 kPa): 187.5 μL acrylamide, 30 μL bisacrylamide, 10 μL APS and 1 μL TEMED. Following TEMED addition, 200 μL of gel solution was immediately pipetted onto a flat Perspex plate and a functionalized coverslip was placed on top. Following polymerization, gels were removed from the Perspex using a square-edged scalpel and hydrated by incubation in PBS for 1-2 hours. Gels were then incubated with cell culture media overnight before use. Where drug treatments were used, these were also included in pre-incubation media. Cells were plated in plastic 12-well plates (Thermo Scientific) 24 hours before a confinement experiment. Where two different cell lines were compared (eg MCF10A and MCF10A+Ras^V12^), both were plated in the same well separated by a PDMS divider that was removed before confinement. This allowed direct comparison of different cell types under the same gel. To confine the cells, the polyacrylamide-coated coverslips were attached to soft PDMS (1:20 cross-linker: base mix) cylinders of approx. 17mm height and 15mm diameter, which were then attached to the lid of the plate, lowered onto the cells and secured with tape. This method of confinement is described in detail by Le Berre et al ^24^. Western Blot Cells were lysed using chilled RIPA buffer on ice. Protein concentration was determined using Bradford reagent and samples were run on a 4-12% Tris/Bis gel (Invitrogen). Protein was then transferred to an Immobilon-P PVDF membrane (Millipore), which was probed using antibodies against the following proteins: phospho-ERK1/2 (Thr202/Tyr204) (Cell Signaling 9101) ERK1/2 (Cell Signaling 9102), phospho-Akt (Ser473) (Cell Signaling 4058), Ras (Cell Signaling 3965), E-cadherin (Invitrogen ECCD2) and GAPDH (Thermo Fisher MA5-15738) and then anti-mouse, anti-rabbit and anti-rat HRP-conjugated secondary antibodies (Dako). ECL detection was carried out using the Immobilon Crescendo HRP substrate (Millpore) in an ImageQuant LAS4000 (GE Healthcare).

### Statistical Analysis

Data analysis was carried out using FIJI and Microsoft Excel. Graphs were produced using Graphpad Prism. Bar charts show mean with error bars showing standard deviation and Box-and-whisker plots use the Tukey method to identify outliers. For live cell imaging, cell shape measurements and gel confinement experiments, graphs show data pooled from at least two independent experiments (normally three). For western blotting and AFM, at least two independent experiments were carried out and one representative dataset is shown. Statistical testing was carried out using Graphpad Prism. Unless otherwise stated, p values were calculated using the Mann-Whitney two-tailed test. *p<0.05 **p<0.01 ***p<0.001.

## Acknowledgements

The authors thank Ewa Zlotek-Zlotkiewicz and Mael Le Berre for help with the gel confinement experiments, Andrew Vaugham, Ki Hng and John Gallagher for microscopy support and Susana Godinho and Julian Downward for reagents. We also thank Susana Godinho, Guillaume Charras, Nitya Ramkumar and Zaw Win for comments on the manuscript.

